# Female C57BL/6J mice perform distinctive urination behaviour accompanied by ultrasonic vocalisations sequences with a stereotypic temporal organisation

**DOI:** 10.1101/2025.02.04.635841

**Authors:** Fabrice de Chaumont, Gaëlle Yvenou, Ana Perez Villalba, Yann Hérault, Thomas Bourgeron, Elodie Ey

## Abstract

Ultrasonic vocalisations (USVs) are largely studied in mice as a marker of social communication. These USVs are usually recorded during short social encounters in unfamiliar test cages. In the present study, we explored how freely interacting pairs of C57BL/6J adult female mice spontaneously use USVs over long-term monitoring. In this situation, we discovered that these mice display a previously undescribed behaviour: they emit specific USV sequences while depositing a large quantity of urine in a corner of the cage. The most striking feature of USVs accompanying this vocalised urination behaviour was the stereotyped duration of the inter-USV intervals. The variability in the frequency of occurrence of this behaviour was important between pairs. Interestingly, when accompanied by the specific USV sequence, urination was correlated with a significant increase in locomotor activity in both the emitter and the cage mate, in contrast with urination without USVs. Altogether, the observation and description of this vocalised urination behaviour argue for exploring mouse vocalisations at the sequence level to understand the USV-behaviours interactions.

## 1. Introduction

Throughout their lives, mice emit ultrasonic vocalisations (USVs). Knowledge about the behavioural contexts of emissions of USVs is currently fragmented. Juvenile and adult mice of both sexes utter these signals when interacting with same-sex conspecifics (D’Amato and Moles, 2001; Panksepp et al., 2007; Chabout et al., 2012; Ferhat et al., 2016) and more rarely when exploring a new environment (Mun et al., 2015) or during restrain stress (Chabout et al., 2012; Lefebvre et al., 2020). Adult males also emit USVs when interacting with females (Holy and Guo, 2005). During short-term same-sex provoked interactions, juvenile and adult male and female mice emit USVs mostly in direct social contact (Chabout et al., 2012; Ey et al., 2018). During spontaneous same-sex long-term recording, females emit most USVs when they are highly aroused in intense social contacts, while males vocalise rarely and mostly when they are not in contact with others (de Chaumont et al., 2021). In addition, adult males emit USVs in close interactions with females, especially before and during mounting or attempts of mounting (Matsumoto and Okanoya, 2021). Altogether, the specific behavioural contexts synchronous with the emissions of USVs has been only partially examined with different precision levels.

At the call level, a striking aspect of mouse USVs is the diversity in the acoustic structure of each call. This has lead scientists to categorise them in 2 to 11 human-based call types (Holy and Guo, 2005; Wang et al., 2008; Enard et al., 2009; Scattoni et al., 2011; Hoffmann et al., 2012; Schmeisser et al., 2012). Other approaches use clustering based on acoustic features to deal with this diversity (Hammerschmidt et al., 2012; Sangiamo et al., 2020). These different call types are assembled in sequences or bursts. Sequences are successive USVs separated by inter-call intervals of less than 230-750 ms, depending on age/sex/context (Holy and Guo, 2005; Ey et al., 2013; de Chaumont et al., 2021). Sequences are separated from one another by inter-sequence intervals longer than these thresholds.

This sequence level was used to study the behavioural effects in recipients of USVs. In pups, sequences of isolation calls trigger retrieval from the mother (Zippelius and Schleidt, 1956; Sewell, 1970). In male-female interactions, the female’s proximity is facilitated by vocalising males (Pomerantz et al., 1983; Hammerschmidt et al., 2009). Female USV sequences triggered only a slight avoidance of the call source by females and subtle increased exploration by males (Wöhr et al., 2011).

In the present study, we therefore advocate to consider USVs at the sequence level to examine the potential functions of USVs during spontaneous emission. We focused on female mice since this sex emitted the highest number of USVs during long-term recordings in a specifically designed environment serving as home cage (**Figure 1A**) (de Chaumont et al., 2021). This approach allowed us to identify a specific behaviour during continuous monitoring of pairs of mice, namely *vocalised urination*: females deposit a large quantity of urine (**Figure 1B**) while uttering specific USV sequences. These sequences made of acoustically simple USVs separated by steady time intervals (**Figure 1C**) were acoustically and rhythmically different from USV sequences given during other behaviours such as the follow behaviour. This vocalised urination behaviour was observed in both mice of each pair, and distributed over day and night times (see one example in **Figure 1D**). To test the potential functions of this specific behaviour, we analysed the spontaneous behaviour of the pairs of females and designed a playback experiment.

**Figure 1:**
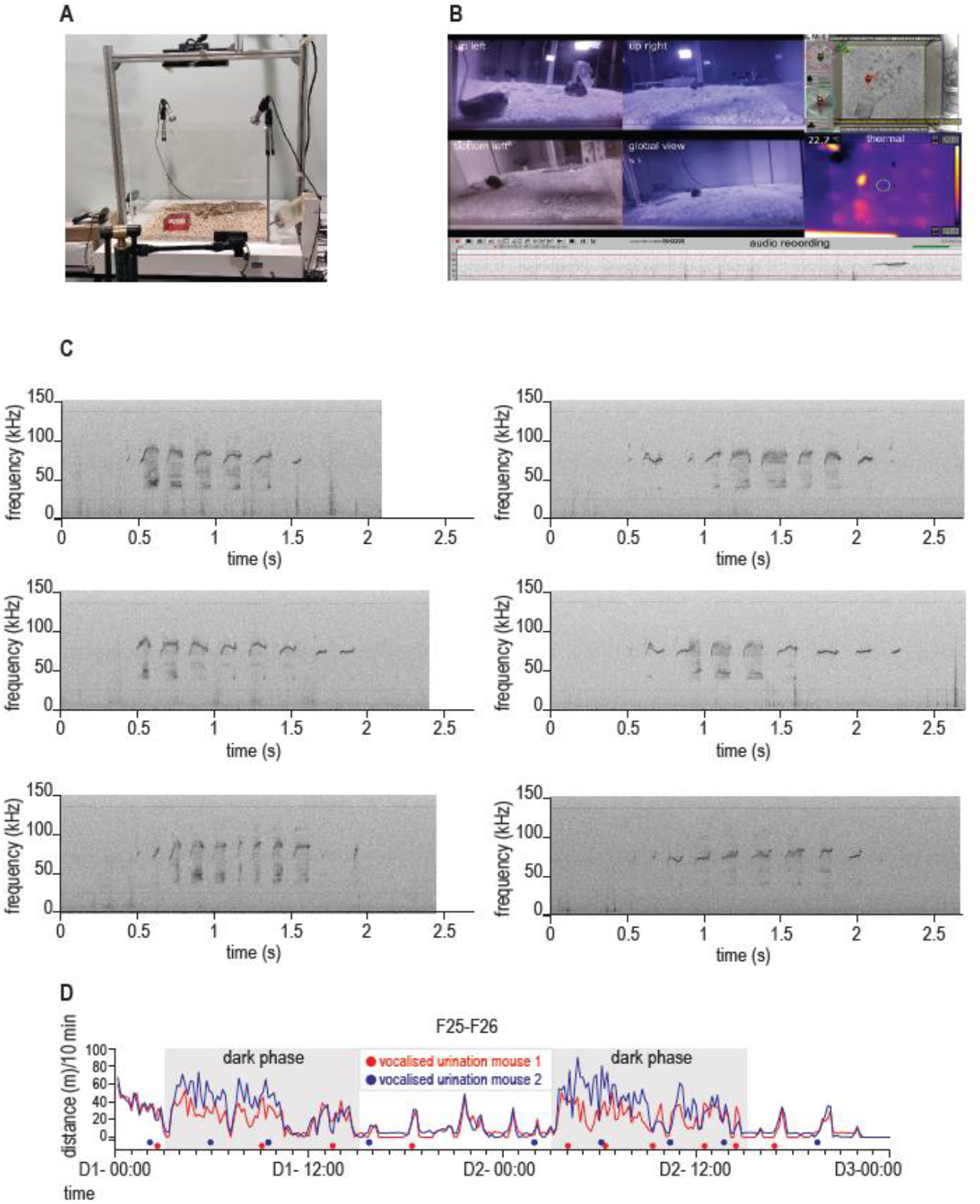
Continuous monitoring reveals a new behaviour combining urination and specific USV sequences in female mice: the *vocalised urination*. A. Experimental set up used to continuously record the behaviours and USV emission of pairs of mice, combining behavioural monitoring (Live Mouse Tracker system), ultrasonic acoustic recordings (Avisoft ultrasonic microphones), thermal camera (on top) and IR cameras on the sides. B. View of the synchronised capture of side views, Live Mouse Tracker tracking, thermal view and audio recordings; on this screenshot, the female tagged in green is urinating in the upper left corner while starting to emit as USV sequence visible on the spectrogram panel at the bottom. C. Spectrograms of six specific USV sequences emitted by the same female while performing the *vocalised urination* behaviour (sampling frequency: 300 kHz, 1024 FFT points; 16-bits format; Hamming window, 75 % overlap). D. Timeline of activity of one pair of females computed per 10 min time bins displaying the distribution of vocalised urination over the two days of recordings.

## 2. Methods

### 2.1. Animals

Cohort 1: 12 C57BL/6J female mice were purchased from Charles River Laboratories (L’Arbresle, France) at 6 weeks of age. Upon arrival, mice were housed in pairs that were conserved throughout the experiment. After one week of acclimation, we identified each individual with a subcutaneous Radio Frequency Identification (RFID) tag (APT12 PIT tags; Biomark, Inc., Boise, The United States of America) inserted under gas anaesthesia (Isoflurane) with local analgesia (Lidor 20 mg/ml, with 40 μl/10 g mouse). RFID tags were located in the lower part of the left flank. Mice were allowed to recover for at least one week.

Cohort 2: We generated a cohort of 12 males and 12 females of C57BL/6J mice in our animal facility. These mice were identified through ear punches (maximum 1 mm diameter) at weaning at 4 weeks of age and housed in same-sex groups of four.

Animals were manipulated using the Plexiglas tunnel placed in their housing cage. Habituation to this manipulation was done at least 3 days before starting the experiments. In compliance with ethical rules and regulatory requirements on use and welfare of laboratory animals, all procedures have been reviewed and approved by our ethical committee (Com’Eth, CE17) registered at the French Ministry of Research under the references APAFIS #15692-2018062715092398 v3 for RFID tag insertion necessary for continuous recordings and APAFIS #42672-2023032116598365 v3 for playback experiments.

### 2.2. Continuous monitoring of pairs of familiar individuals

We continuously recorded dyadic interactions between two familiar female mice of the same age (8-12 weeks of age) from cohort 1. Altogether, we recorded six pairs of females. Recordings started between 03:00 and 04:00 PM and lasted undisturbed 47h long, that is, two days and nights minus 1h for cleaning in between two recording sessions. We coupled the Live Mouse Tracker (LMT) system (version 931; (de Chaumont et al., 2019)) with the Avisoft Ultrasound Gate 416 (300 kHz sampling rate, 16-bit format; trigger: level of this channel; pre-trigger: 1 s; hold time: 1 s; duration > 0.005 s; trigger event: 2 % energy in 25-125 kHz with entropy < 50%; Avisoft Bioacoustics, Glienecke, Germany) connected to a CM16/CMPA microphone (Avisoft Bioacoustics, Glienecke, Germany) (**Figure 1A**). LMT and Avisoft systems were synchronised based on the protocol described in (de Chaumont et al., 2021). We also added the synchronised capture of thermal data using a thermal camera (Flir C3) and side views using webcam (Logitech C920 HD Pro with removed Infra Red filter) and Infra Red projector positioned at the sides of the LMT cage (**Figure 1B**). We manually checked the probability of the identity of the USV emitter by comparing the amplitude between the two channels given the orientation of the animals.

Behavioural data were explored using the analysis scripts provided on github (link to be added upon publication) and the Live Data Player from LMT. USVs were detected and analysed automatically using LMT USV Toolbox (de Chaumont et al., 2021). An online version is available on https://usv.pasteur.cloud (offline version downloadable at the same location too).

### 2.3. Playback experiment

Mice from cohort 2 underwent the playback experiment at 6 weeks of age in a sound-attenuated chamber between 9:00 and 12:00 AM. The setup was constituted by an elevated platform of 50 x 20 cm, with a loudspeaker located at each extremity 8 cm away from the end of the platform. Microphones were placed 25 cm above each extremity of the platform to record the broadcasted signals and potential USVs from the tested animals.

The audio files used for the playback experiments were 2-channels files, with background noise on one channel and a USV sequence (=signal) on the other channel. As signals, we used two USV sequences recorded in one pair of the cohort 1 (unfamiliar to the mice of cohort 2). One USV sequence occurred during vocalised urination and the other USV sequence occurred during long chase (i.e., long and intense following) between the two females of the pair. We selected two background noise sequences (without USVs) of the same duration as the signal sequences. Each audio file consisted in two repetitions of the same signal + background noise sequence separated by 2 s of silence. All recordings were high-pass filtered at 30 kHz. Our playbacks were broadcasted at 50-70 dB at 20 cm (after calibration of the microphone). The side of the signal and the order of presentation of the urination or long chase USV sequence over playback 1 and playback 2 were balanced over all animals.

Day 1 consisted in the habituation to the testing setup and to the temporal sequence of events that occurred the next day for the playback test. For that purpose, we released each mouse on the platform and let it explore for 1 min. After 1 min, we constrained the animal under a small enclosure (8 x 8 x10 cm; grid sides facing each loudspeaker) in the middle of the platform for 10 s. Then, we lifted the grid enclosure and let the mouse explore the platform for 2 min. We then constrained again the animal under the grid enclosure in the middle of the platform for 10 s. After that time, we lifted the grid enclosure and let the mouse explore again the platform for 2 min. We put the animal back in its housing cage after this habituation.

On day 2, we conducted the playback test. As in the habituation the day before, we placed the mouse on the platform and let it explore for 1 min. After 1 min, we constrained the animal under the grid enclosure in the middle of the platform and played back the audio file with the USV sequence synchronous with urination or with long chase behaviour as signal versus background noise (playback 1). We lifted the grid enclosure right after the first occurrence within the audio file. Therefore, the second repetition occurred right after lifting the grid enclosure, while the animal was already free to explore. We let the animal explore the platform for 2 min and we constrained again the animal under the grid enclosure in the middle of the platform for 10 s. We then played back the audio file with the other signal on the other side compared to the first playback (playback 2). As in playback 1, we lifted the grid enclosure right after the first occurrence within the audio file. Therefore, the repetition occurred right after lifting the grid enclosure. We let the animal explore the platform for 2 min. At the end of this exploration time, we put the mouse back in its housing cage.

### 2.4. Analyses

All analyses were run in Python. LMT analyses used the available scripts on GitHub (https://github.com/fdechaumont/lmt-analysis). Other scripts will be available upon request.

#### 2.4.1. LMT recordings

We used a linear mixed model (LMM) with the behavioural context as a fixed factor and the pair of mice as a random factor to compare the features of the USV sequences or of the USVs between vocalised urination events and other behavioural contexts. We applied a Bonferroni correction for multiple testing when comparing the USV sequences / USVs given during vocalised urination versus all other contexts (five different contexts). We did not compare the other five contexts within each other since we were only interested in the specificities of USV sequences given during vocalised urination compared to all other contexts. We used the python script *ExploreUSVTimeLineActivity.py*.

Timelines were generated in python with the script *ExploreUSVTimeLineActivity.py*. The effects of vocalised urination or urination without USVs (silent urination) on the behaviour of the mouse urinating and the cage mate were computed using the *exploreUrinationEvents.py* script. To test the effect on the activity, we used a one-sample t-test to compare the difference in activity before and after the urination event (delta distance) with the reference value (0). The durations of vocalised urination and silent urination events were compared using LMM with context as a fixed factor and individual as a random factor in the script *exploreUrinationEvents.py*.

#### 2.4.2. Playback test

In the playback experiment, we manually scored the initial reaction of the mouse to the different acoustic stimuli. We scored whether the animals oriented or not (attraction, repulsion or no reaction) toward the signal. We compared the scores between the playback of USVs synchronous with vocalised urination events and the playback of USVs synchronous with long chase behaviours using a non-parametric paired Wilcoxon test given the small sample size. Using EZTrack (Pennington et al., 2019), we measured the distance travelled by the animals following each playback. We compared the distances travelled in the two minutes of free exploration after the playback of USVs synchronous with urination behaviour and after the playback of USVs synchronous with long chase behaviour using a non-parametric paired Wilcoxon test. We used an ethological keyboard (BORIS (Friard and Gamba, 2016)) to manually annotate on the videos the time spent in each zone of interest. We defined the proximity zones as the 10 cm of the platform the closest to the loudspeakers. We compared the proportion of time spent in the proximity zone of the signal (either USVs synchronous with urination behaviour or USVs synchronous with long chase behaviour) (over the total time spent in both proximity zones) to a chance value of 0.5 (meaning no preference for any proximity zone) using a one-sample non-parametric Wilcoxon test.

## 3. Results

### 3.1. Female mice perform a specific behaviour combining urination and USVs: vocalised urination

When manually exploring the 2-days recordings of spontaneous USVs of six pairs of adult female C57BL6J mice, we identified 90 occurrences of a previously undescribed behaviour combining micturition and USVs. These specific behavioural events that we named *vocalised urination* events consisted in a sequence of highly reproducible and stereotypic behaviours. They started with one of the pair of mice sitting in a corner (one of the two corners the furthest away from the nest), facing the centre of the cage and bouncing back its tail along the wall (**Figure 1B**, up left view). The female emitted a stereotyped USV sequence (see examples in **Figure 1C**) while depositing urine. The micturition was confirmed on synchronous side views and thermal recordings (**Figure 1B**). We observed that vocalised urination events were associated with large deposits of urine, in contrast with the small quantities that female mice usually drop while moving (Coquelin, 1992).

### 3.2. Vocalised urination is spontaneous and follows an unpredictable pattern

Vocalised urination events integrate two very different stimuli: acoustic signals through USVs and chemical signals in urine. In order to understand the relationship between these two types of signals, we quantified and analysed single and overlapping production of both signals for two consecutive days and nights of recordings. Every pair of females performed between 0 and 12 vocalised urination events (mean+/-standard deviation = 7.5+/-3.9 sequences per individual; **Figure 1D & 2A-E**). Vocalised urination events occurred both during days and nights. In addition, the same behaviour of urination but without USVs (that we named *silent urination* events) was displayed by the females up to 9 times (mean+/-standard deviation = 2.5+/-3.1) over the two days and nights (in total 28 silent urinations were recorded). These silent urination events (8.0+/-0.6 s, mean+/-SEM) were significantly shorter than vocalised urination events (10.1+/-0.6 s, mean+/-SEM; LMM: log-likelihood=-344.8, beta=3.34, SE=1.203, p=0.005). As a comparison, each female mouse performed 42.4+/-5.4 long chases accompanied by USVs lasting 13.3+/-0.3 s (mean+/-SEM), and 5.6+/-0.7 silent long chases lasting 11.1+/-0.9 s over the two days of recordings, confirming the previously described increased duration of behavioural events accompanied by USVs in comparison with similar behaviours without USVs (de Chaumont et al., 2021).

There was a large variability in the number of vocalised urination events versus silent urination events between mouse pairs. For instance, the pair F25-F26 showed a high number of vocalised urinations (**Figure 1D**), while the pair F27-F28 performed few vocalised urinations and a majority of silent urinations (**Figure 2C**). Among the 118 urination events (vocalised and silent) recorded, 40 occurred within one hour of another one, showing no specific temporal association. Altogether, female mice did not follow a specific temporal pattern for vocalised urination events, that were considered as spontaneous and unpredictable given the current data.

**Figure 2:**
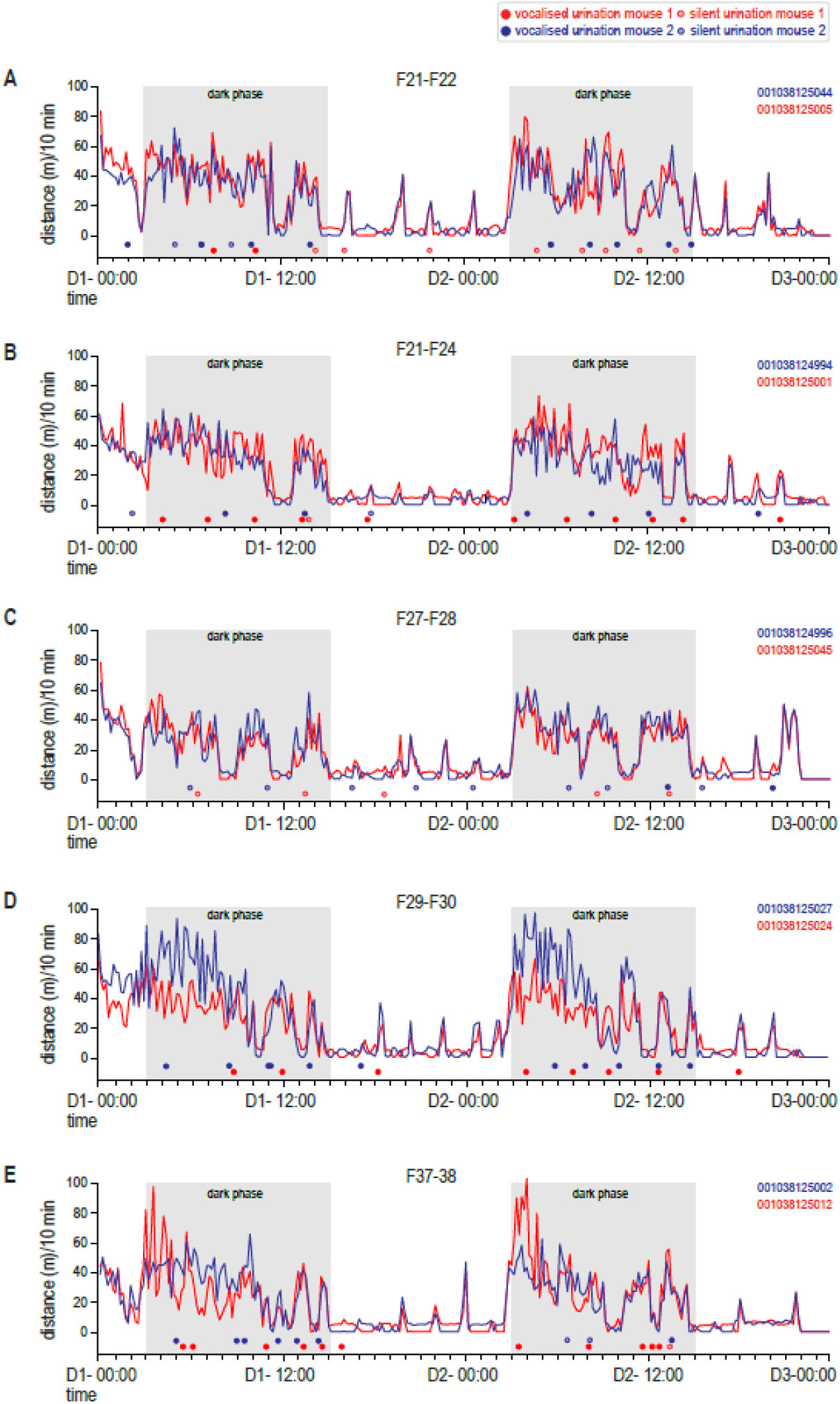
Urination events with or without USVs according to the activity period. Each panel (A-E) represents the distribution of vocalised urination and silent urination events as well as the activity per ten minutes time bin for a pair of females over two days and two nights.

### 3.3. USV sequences emitted during vocalised urination events present with stereotypic temporal organisation of acoustically simple USVs

We compared temporal and acoustic features of vocalised urination events with other behavioural events such as long chase. Vocalised urination events lasted 10.1+/-0.6 s (mean +/-SEM) and were significantly longer in comparison with events such as follow, train2, oral-genital contacts, nose-nose contacts and approach before a contact (**Figure 3A**). Vocalised urination events were the closest (in duration) with long chase events, but still significantly shorter (**Figure 3A**).

**Figure 3:**
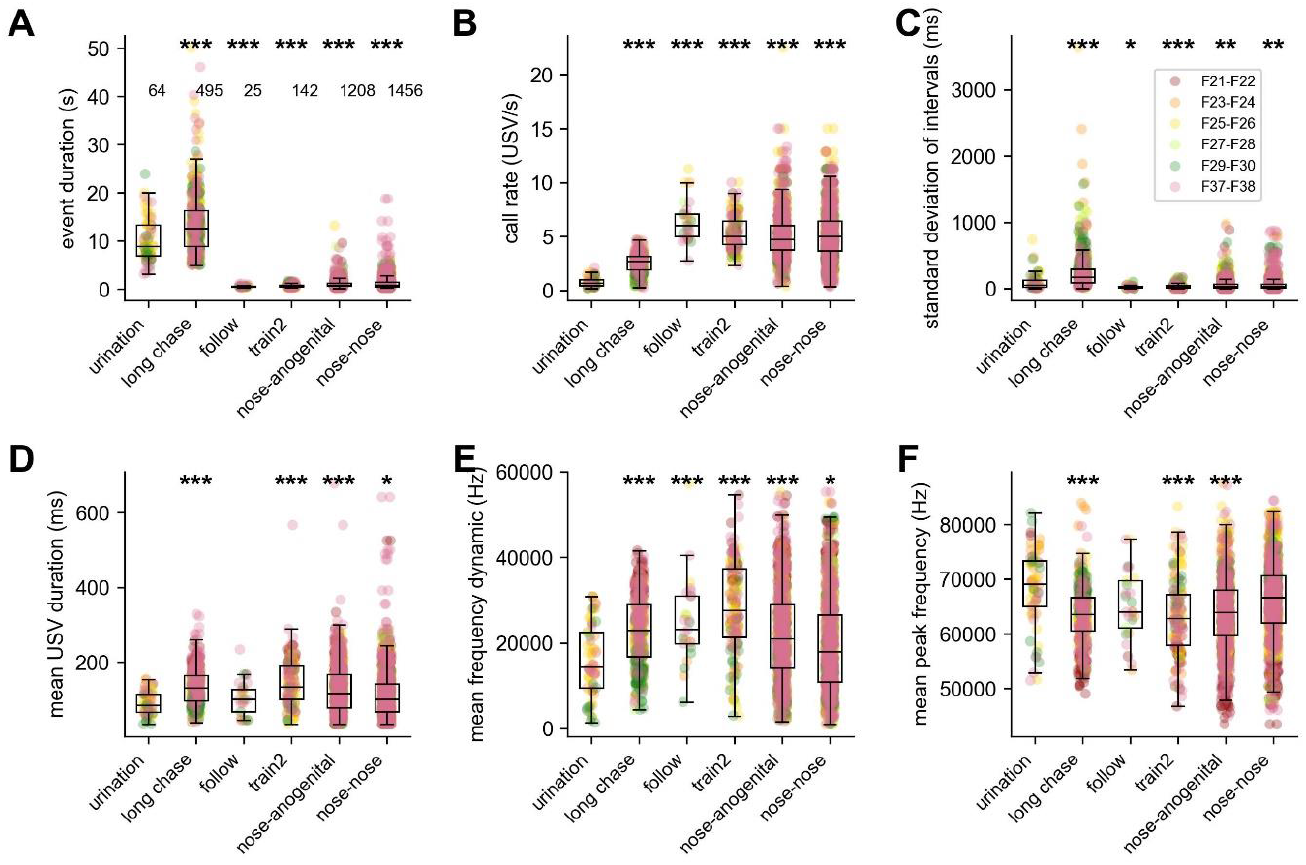
Temporal organisation and acoustic features of USVs synchronous with vocalised urination events compared to other behavioural contexts. A. Duration of the behavioural events that occurred simultaneously with USVs (vocalised urination events versus all others such as long chase, follow, train2, nose-anogenital and nose-nose contacts). B. Rate of emission within a USV sequence according to the behavioural contexts. C. Standard deviation of the duration of intervals between USVs within a USV sequence according to the simultaneous behavioural contexts. D. Mean duration of the USVs for each sequence according to the simultaneous behavioural contexts. E. Mean range of frequency covered within a USV sequence according to the simultaneous behavioural contexts. F. Mean peak frequency within a USV sequence according to the simultaneous behavioural contexts. The numbers of behavioural events examined figure on panel A. Linear mixed model with behavioural context as a fixed factor and pair as a random factor to compare vocalised urination versus each of the other contexts (long chase, follow, train2, nose-anogenital contacts and nose-nose contacts). Colours indicate the pair of mice in which USVs were recorded. Stars represent statistical significance, after correction for multiple testing: ^*^: p<0.01, ^**^: p<0.002, ^***^: p<0.0002.

We quantified the specificities of the temporal organisation and the acoustic characteristics of USVs emitted during vocalised urination (see examples of spectrograms of USVs emitted during vocalised urination in one pair of females in **Figure 1**). The temporal organisation of USVs simultaneous with vocalised urination events differed in comparison with USVs during other behavioural contexts. The call rate (USVs per second) within the events was significantly lower during vocalised urination events than during all other types of behaviours (**Figure 3B**). The most striking feature was that the intervals between USVs were more stereotyped during vocalised urination events than during other behaviours such as long chase, with a significantly lower standard deviation of interval durations in USVs synchronous with vocalised urination (**Figure 3C**).

Next, we characterised the acoustic structure of USVs associated with vocalised urination. These USVs had a significantly shorter mean duration compared to USVs given during long chase, train2, nose-anogenital contacts and nose-nose contacts, but were similar to USVs given during follow events (**Figure 3D**). USVs synchronous with vocalised urination events were also less frequency-modulated compared to USVs given during long chase, follow, train2, nose-anogenital and nose-nose contacts (**Figure 3E**). The mean peak frequency of USVs during vocalised urination events was similar with those given in follow behaviour, while it was significantly higher compared to those given in long chase, train2 and nose-anogenital contact (**Figure 3F**). Altogether, USVs during vocalised urination events were acoustically simpler than USVs given in other behavioural contexts.

### 3.4. Vocalised urination events triggered an increased activity but no approach of the conspecific

Vocalised urination is a complex behavioural pattern that could have an effect in the mouse performing the behaviour or in the cage mate. As a starting point, we examined in which contexts vocalised urination occurred and whether these contexts were perturbed. For that purpose, we first evaluated whether the vocalised urination events occurred during specific behaviours of the cage mate. In the observation of 90 vocalised urination events, the cage mate was idling in the nest (33/90), idling somewhere else in the cage (16/90), foraging (16/90), or moving alone (25/90). Among these 90 events, only one started with the two animals in contact and only 13 included an approach before a contact. These observations suggested that vocalised urination events were not triggered by or inducing closeness, contact or repulsion between the mouse performing the behaviour and the cage mate.

Then, we tracked locomotor activity before and after vocalised urination events to evaluate possible changes associated with this behaviour. For that purpose, we compared the distance travelled before and after urination for the mouse that urinated and for the cage mate during different time windows (between 1 min and 20 min of duration). Vocalised urination events were followed by a significant increase in the locomotor activity of both animals between 2 and 10 min after vocalised urination, and up to 15 min in the mouse that performed the vocalised urination (**Figure 4A & B**, data in grey). As a comparison, we conducted the same analysis before and after silent urination events to evaluate whether depositing urine alone (silent urination) had a similar effect on the activity of any of the paired animals. In contrast to vocalised urinations, none of the animals (urinating mouse or cage mate) showed any significant change in the activity before or after silent urination (**Figure 4A & B**, data in yellow). These data suggested a correlation between the vocalised urination and locomotion upheaval that is not reproduced by the urination alone (silent urination). The specific USVs emitted during vocalised urination events, complemented with motor behaviour and urination, would have an arousing effect in adult mice. Interestingly this effect was shared by the urinating mouse and the cage mate, which could mean that the increased activity starting first in the mouse performing the vocalised urination behaviour is affecting her cage mate. Interestingly, when computing the difference of locomotor activity before and after a long chase with or without USVs, we did not observe a similar pattern. Indeed, the increased activity was clearly visible in both mice of the pair at any time points (**Supplementary Figure S1**), suggesting that in long chases the emission of USVs is reflecting a high arousal of both mice.

**Figure 4:**
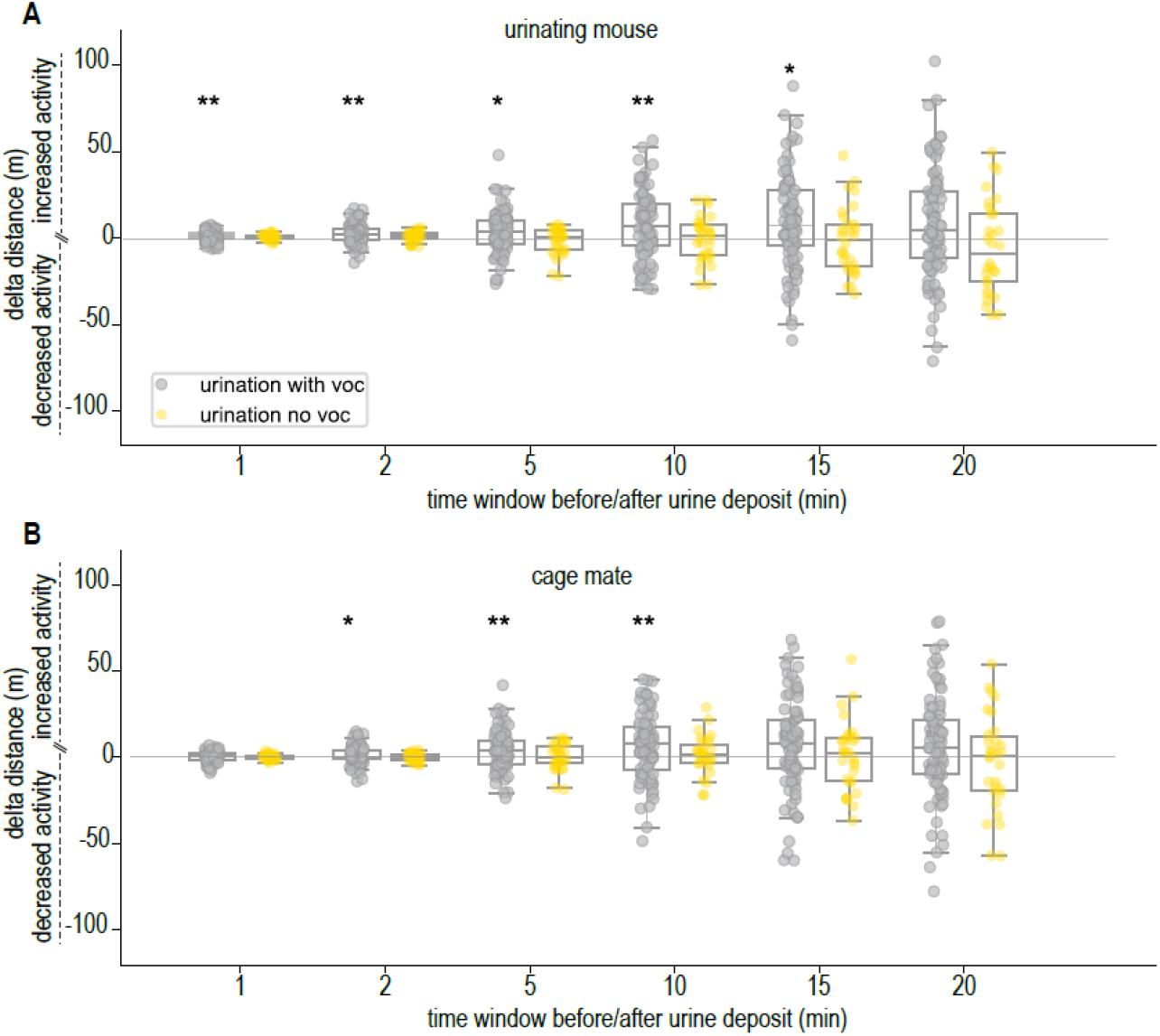
Behavioural effects of vocalised urination and silent urination events. A. Difference in the distance travelled by the urinating mouse before and after the micturition with USVs (vocalised urination; data in grey) or without USVs (silent urination; data in yellow) according to the time window. B. Difference in the distance travelled by the cage mate before and after the micturition with USVs (vocalised urination; data in grey) or without USVs (silent urination; data in yellow) according to the time window. We used one-sample t-tests to compare to zero the difference of the activity before/after a behavioural event (activity after minus activity before; therefore it is positive it means that the activity increases after urination, while if it is negative it means that the activity decreases after urination). ^*^: p<0.05, ^**^: p<0.01.

To test whether this increase of activity was related to a visit of the urination corner by the cage mate, we compared the probability to visit the corner after vocalised urination and after silent urination. This probability of visits could be computed in 84 vocalised urination events and in 30 silent urination events. We chose a time window of 2 min as it is the first time-window in which the activity of the cage mate was significantly increased (**Figure 4B**). Under these conditions, vocalised urinations were followed by a visit of the corner by the cage mate in 58% of cases (49/84), while silent urinations were followed by a visit of the corner by the cage mate in 57% of cases (17/30). As a comparison, a stay in the corner without urination (but with a duration that could allow urination) was followed by a visit of the cage mate in 59% of cases (2126/3579). Therefore, the presence or absence of USVs during micturition did not influence significantly the probability to visit the corner by the cage mate. This suggested that vocalised urination events do not specifically attract or repel conspecifics.

### 3.5. Playback of USVs recorded during female vocalised urinations had opposite effects in female and male mice

In order to test further this assumption in a more controlled setting, we examined the behavioural effect of the acoustic signals alone (i.e., the olfactory cues were not present) in the vocalised urination events in both sexes. Using a specifically designed playback platform (**Figure 5A**), we compared the behavioural reactions of male and female mice when hearing one of these specific USV sequences emitted during vocalised urination events (**Figure 5B**) versus one USV sequence recorded during long chase events as control stimulus (**Figure 5C**). We played back the two types of USV sequences from C57BL/6J females to unfamiliar C57BL/6J males and females aged of 2 months (**Figure 5D**). Mice of both sexes reacted to playback of both types of USV sequences by freezing or orienting. This showed that they perceived the signals.

**Figure 5:**
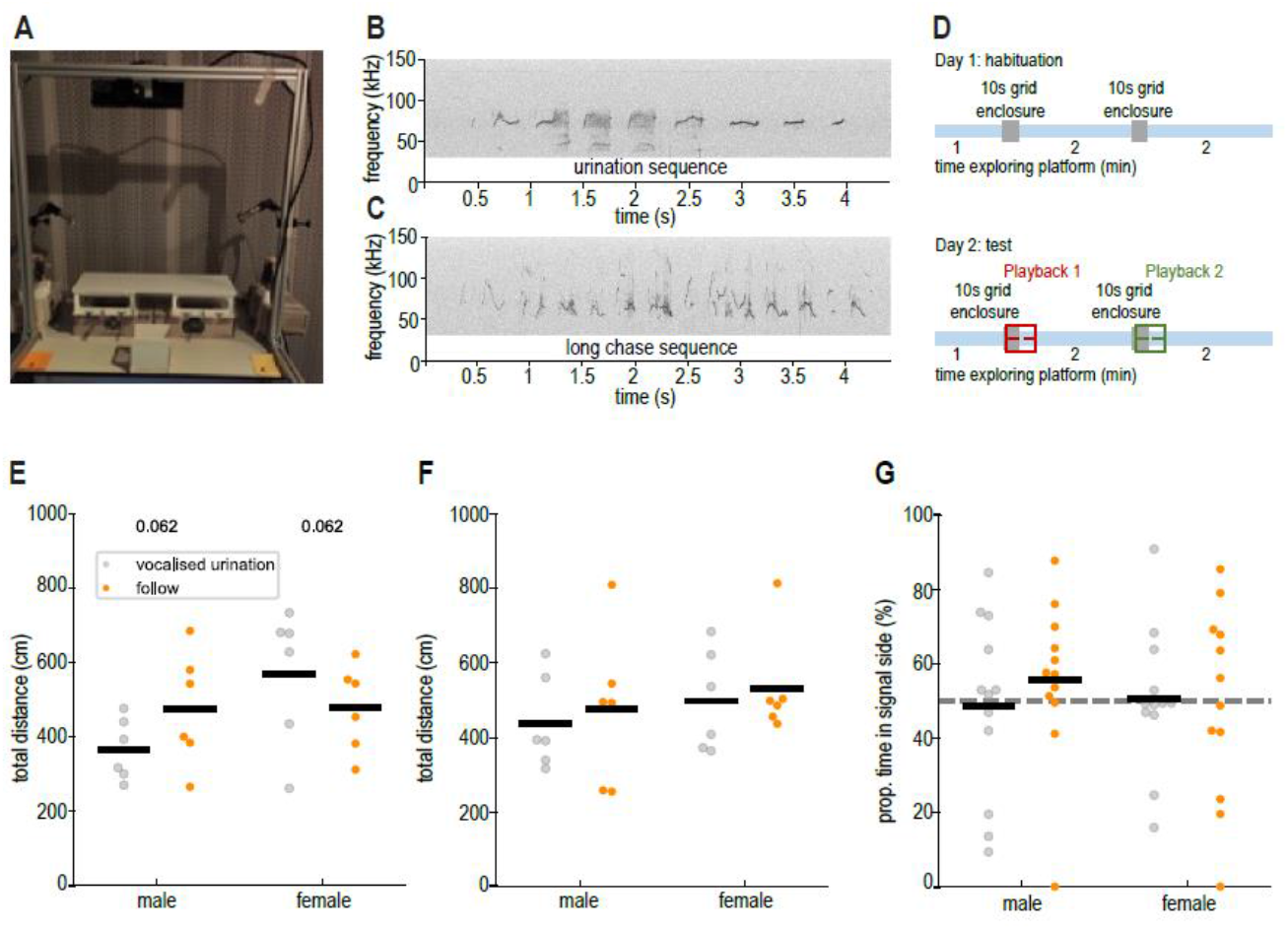
Activity and behavioural reaction to the playback of USV sequences recorded during vocalised urination versus USVs recorded during long chase behaviour. A. Playback setup used to test the behavioural reaction to playback of USVs synchronous with vocalised urination versus background noise and of USVs synchronous with long chase behaviour versus background noise. The grid enclosure is lying on the table in the foreground down to the platform in the middle of the setup. B. Spectrogram of a USV sequence recorded during vocalised urination, including 9 USVs (300 kHz sampling frequency; 1024 FFT points, 16 bits, Hamming window, 75 % overlap). C. Spectrogram of a USV sequence recorded during long chase behaviour including 13 USVs (300 kHz sampling frequency; 1024 FFT points, 16 bits, Hamming window, 75 % overlap). D. Protocol used to habituate the mice to the playback setup (upper panel: Day 1) and to conduct the playback experiment (lower panel: Day 2). E. Distance travelled in the two minutes following the playback of USVs synchronous with vocalised urination versus background noise and of USVs synchronous with long chase behaviour versus background noise when urination sequence was presented first. F. Distance travelled in the two minutes following the playback of USVs synchronous with vocalised urination versus background noise and of USVs synchronous with long chase behaviour versus background noise when follow sequence was presented first. G. Proportion of time spent in the proximity zone of the tested signal in the two minutes following the playback of urination USV sequences versus background noise and of long chase USV sequences versus background noise. Numbers represent the p-value of Wilcoxon tests when smaller than 0.1.

Based on our observation of the spontaneous behaviours (increasing activity, **Figure 4**), we expected an increase of the locomotor activity after the broadcast of USVs accompanying vocalised urination events compared to the broadcast of USV sequences synchronous with long chase behaviour in female mice. To test this hypothesis, we measured the total activity over the two minutes after playback. We observed two different behavioural reactions in male and female subjects with two opposite tendencies and these effects were present only when testing USVs given during vocalised urination events first. Indeed, in this condition, females tended to increase their locomotor activity after hearing USVs related to vocalised urination events in comparison with USVs related to long chase events (**Figure 5E**). In contrast, males displayed reduced locomotor activity after hearing USVs synchronous with vocalised urination events in comparison with USVs synchronous with long chase events (**Figure 5E**). This suggests that the previously shown arousal effect of vocalised urination events might be sex-specific. No trends were visible when long chase sequences were tested first (**Figure 5F**) which might be related to the rapid habituation to USV stimuli that has been previously highlighted in playback experiments (Hammerschmidt et al., 2009; Wöhr et al., 2011). After observing these sex-specific reactions, we measured the proportion of time spent in each proximity zone around the loudspeakers to detect any attraction or repulsion toward the acoustic signal tested. Both males and females did not show any significant preference toward the side broadcasting the USV sequences (either synchronous with vocalised urination or synchronous with long chase behaviour), no matter the order of presentation of the signals (**Figure 5G**). As none of the signals triggered place preference, we could not qualify the USV sequences emitted during vocalised urination as a reinforcing or attractive signal per se.

## 4. Discussion

In the present study, we describe a new behaviour in C57BL/6J females, namely vocalised urination: micturition accompanied by specific USV sequences. The USV sequences were characterised by acoustically simple USVs separated by stereotyped inter-call intervals. Vocalised urination was associated with increased activity in both the urinating mouse and the cage mate, that was not found when only urination occurred (silent urination). Playback experiments confirmed the increased activity triggered by USV sequence emitted during vocalised urination in females but not in males. In addition, we did not highlight significant preference for USV sequences emitted during vocalised urination over long chase USV sequences neither in males nor in females.

In a previous study, we recorded pairs of C57BL/6J mice in similar conditions (de Chaumont et al., 2021). Digging back into the data, we found similar vocalised urination events that were performed by males (paired with another male) and in other pairs of females (**Table I**). In this study, we recorded the same animals at the juvenile (5 weeks), young adult (3 months) and older adult (7 months) stages. We observed vocalised urinations at the three stages recorded in both sexes, but there was a large inter-individual variability in the expression of this behaviour, with some animals never performing it and other performing it more than 20 times over three days (**Table I**). This vocal behaviour present in both sexes and already present at the juvenile stage in C57BL/6J mice differs from scent marking, that is used to transmit information even when the emitter is absent (Arakawa et al., 2008). The specificity of vocalised urinations lays in the combination of immediate communication through acoustic signalling, and of long-lasting signals through micturition.

**Table I:** Number of vocalised urination events that were recorded over three days of continuous monitoring of pairs of C57BL/6J mice of 5 weeks, 3 months and 7 months of age. Silent urinations could not be counted since we could not ascertain that the animals were indeed urinating while in the corner (we did not set up side views on this setup).

| Sex | Individual | Pair | 5 weeks | 3 months | 7 months |
| --- | --- | --- | --- | --- | --- |
| males | M40 | M40_M41 | 0 | 0 | 0 |
| males | M41 | M40_M41 | 15 | 11 | 7 |
| males | M42 | M42_M43 | 0 | 0 | 0 |
| males | M43 | M42_M43 | 3 | 0 | 0 |
| males | M44 | M44_M45 | 2 | 0 | 0 |
| males | M45 | M44_M45 | 5 | 0 | 0 |
| males | M46 | M46_M47 | 0 | 1 | 0 |
| males | M47 | M46_M47 | 0 | 0 | 0 |
| females | F13 | F13_F14 | 21 | 15 | 22 |
| females | F14 | F13_F14 | 19 | 12 | 3 |
| females | F15 | F15_F16 | 18 | 13 | 9 |
| females | F16 | F15_F16 | 0 | 1 | 0 |
| females | F17 | F17_F18 | 3 | 10 | 17 |
| females | F18 | F17_F18 | 6 | 11 | 18 |
| females | F19 | F19_F20 | 15 | 20 | 14 |
| females | F20 | F19_F20 | 1 | 13 | 16 |

In another study, recordings of the female progeny of wild house mice *Mus musculus domesticus* also highlighted that females emitted undirected USVs from a corner of their enclosure where they often urinated, not in visual/tactile contact with any other mouse (Hoier et al., 2016). However, as the analysis was run only at the vocalisation level and not at the sequence level, we cannot know whether these USVs were organised in specific sequences. The acoustic characteristics of these USVs suggested that they were generally more complex than the USVs we describe here. Hoier and colleagues suggested that these USVs might be involved in the transmission of reproductive status or territoriality (Hoier et al., 2016). However, in our case, the playback experiment does not support the hypothesis of a robust transmission of the reproductive status of females in C57BL/6J mice. Indeed, in our experiment, males tend to eavesdrop on female USVs emitted during long chase between females (slightly longer time spent in the signal side than in the background noise side) but not on USVs recorded during vocalised urination (**Figure 5G**).

Concerning the potential functions of vocalised urination, we first thought that it could be a mark of dominance. Nevertheless, both mice of the pair performed this behaviour, and not specifically in the centre of the cage (Drickamer, 2001). In males, urine from dominant mice is aversive, while urine from subordinate mice is not (Jones and Nowell, 1989). In addition, the most dominant male house mice excreted larger quantities of urine compared to subordinate ones (Drickamer, 1995). In our data, this was not clear in females and remains to be investigated further at the individual level.

The vocalised urination behaviour that we described here did not seem to function as female enurination (i.e., spraying urine) by female maras *Dolochotis patagonum* that use this behaviour to deter a conspecific (Ottway et al., 2005). Our results suggest that the emission of a specific USV sequence during urination did not trigger a specific approach or retreat behaviour from surrounding conspecifics. At least, these vocalised urination events appeared to induce an increased arousal, since the activity level increased after their emission / perception.

Studying USV functions at the USV sequence level is ethologically relevant, since the specific behavioural context are as likely to be attached to sequences as to the short acoustic signals separately. Here, the stereotyped temporal organisation of the simple USVs during vocalised urination was the most striking feature. This temporal organisation has already been found to be highly relevant for mothers listening to pup isolation calls and for females listening to male courtship USVs (Agarwalla et al., 2023; Perrodin et al., 2023). Therefore, for further analyses in the functions of USVs, we should also look for specific sequences that distinguish from each other in their temporal organisation.

Altogether, monitoring mice continuously allowed the identification of a new behaviour in mice, that could be included in the behavioural repertoire scored in mouse models for different conditions. The ability of following urination events may also be of interest while monitoring kidney or diabetic disease models. For the moment, the significance of the vocalised urination behaviour remains unclear. It might well be a behaviour specific to some strains of mice, as a deviation through inbreeding (Egnor and Seagraves, 2016). Indeed, it was not observed in a mixed background C57BL/6N with sighted C3H/HeH (E. Ey, personal observation). Further studies will examine other strains of mice. These studies should also focus on diversifying the contexts of recording: larger groups of females and males in which hierarchy will be measured as well as pairs and groups of mice from different strains. We should also check whether this behaviour occurs in mice housed singly (a previous study showed that the quantity of urine excreted did not vary with the density in the housing cage in male or female house mice (Drickamer, 1995)) and check the sexual status since female house mice in oestrus produce more urine than females in dioestrus (Drickamer, 1995) to explain inter-pair variations. For all these studies, adapting sound source localisation methods (Matsumoto et al., 2022; Sterling et al., 2023) will be necessary, especially for groups larger than pairs.

## 5. Acknowledgments

The authors would like to thank the members of the research group for their fruitful discussions, and members of the IGBMC and of the ICS in the animal facility for technical support.

This work was supported by the National Centre for Scientific Research (CNRS), the French National Institute of Health and Medical Research (INSERM), the University of Strasbourg (Unistra), the French government funds through the “Agence Nationale de la Recherche” in the framework of the Investissements d’Avenir Program [ANR-10-INBS-07 PHENOMIN to YH] and Investissements d’Avenir labeled IdEx Unistra [ANR-10-IDEX-0002 to YH], a SFRI-STRAT’US project [ANR 20-SFRI-0012 to YH], EUR IMCBio [ANR-17-EURE-0023 to YH], STRAS&ND [to EE and YH], INBS PHENOMIN [ANR-10-INBS-07 PHENOMIN to YH]. APV’s work was supported by Program for the promotion of scientific research, technological development and innovation in the Valencian Community (Spain) CIBEST 2024-114. The funders had no role in the study design, data collection and analysis, decision to publish, or preparation of the manuscript.

