## Supplementary Figure S1 for "Female C57BL/6J mice perform distinctive urination behaviour accompanied by ultrasonic vocalisations sequences with a stereotypic temporal organisation"

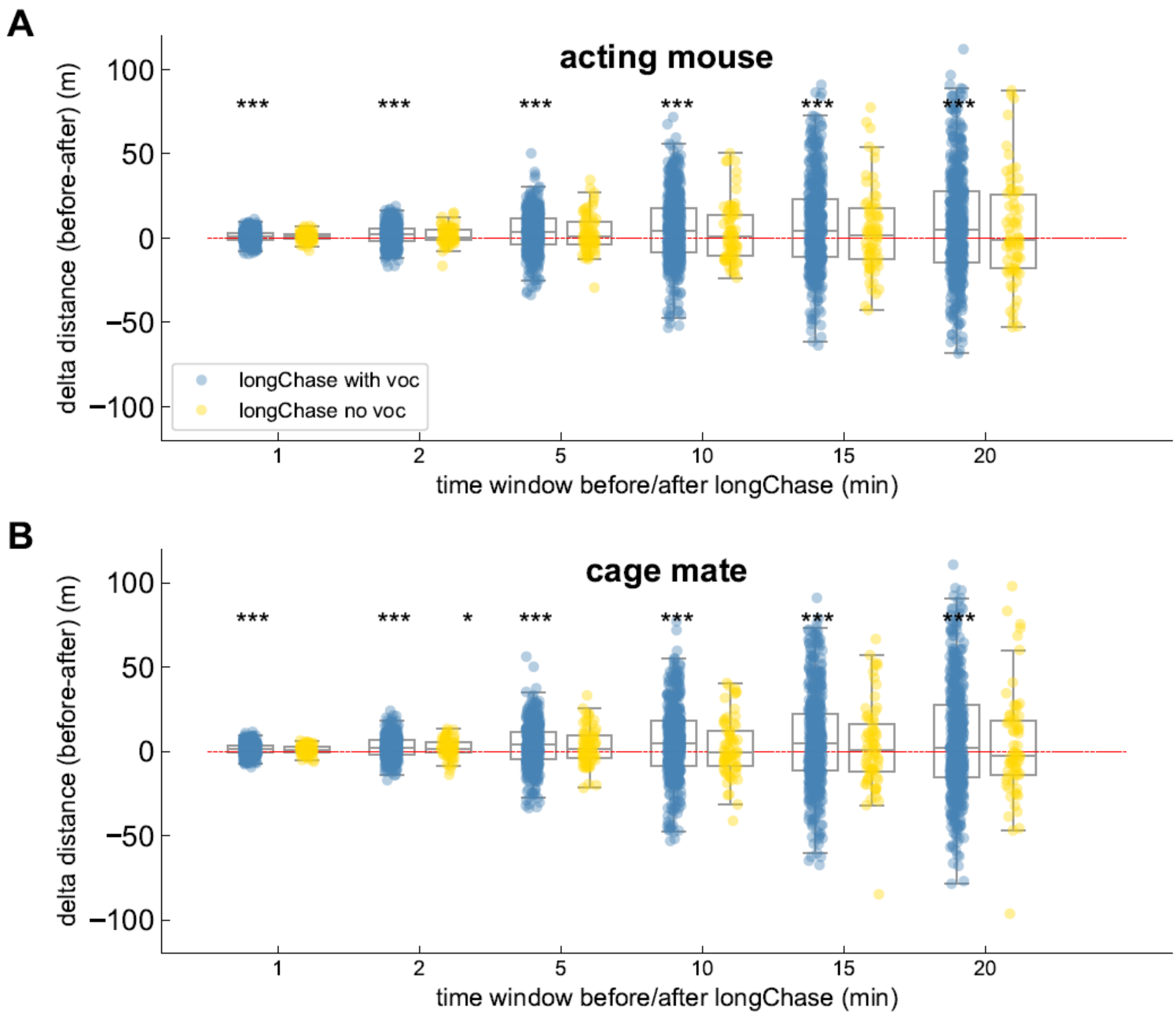


**Supplementary Figure S1: Behavioural effects of vocalised long chases and silent long chases events**. A. Difference in the distance travelled by the chasing mouse before and after the long chases with USVs (vocalised long chases; data in blue) or without USVs (silent long chases; data in yellow) according to the time window. B. Difference in the distance travelled by the cage mate before and after the long chases with USVs (vocalised long chases; data in blue) or without USVs (silent long chases; data in yellow) according to the time window. We used one-sample t-tests to compare to zero the difference of the activity before/after a behavioural event (activity after minus activity before; therefore if it is positive it means that the activity increases after long chase, while if it is negative it means that the activity decreases after long chase). *: $p<0.05$, **: $p<0.01$.
